# Species abundance information improves sequence taxonomy classification accuracy

**DOI:** 10.1101/406611

**Authors:** Benjamin D. Kaehler, Nicholas A. Bokulich, Daniel McDonald, Rob Knight, J. Gregory Caporaso, Gavin A. Huttley

## Abstract

Popular naive Bayes taxonomic classifiers for amplicon sequences assume that all species in the reference database are equally likely to be observed. We demonstrate that classification accuracy degrades linearly with the degree to which that assumption is violated, and in practice it is always violated. By incorporating environment-specific taxonomic abundance information, we demonstrate that species-level resolution is attainable.

## Main

Advances in high-throughput DNA sequencing and bioinformatics analyses have illuminated the crucial roles of microbial communities in human populations and planetary health^12^ and enable microbiome meta-analysis on a massive scale^3^. An important step in characterizing microbial communities is classification of short marker-gene DNA sequences (e.g., bacterial 16S rRNA genes) to infer taxonomic composition.

Short marker-gene sequence reads often contain insufficient information to differentiate species using conventional methods^4–8^. However, current best practices rely on species-level classification to circumvent well-documented inconsistencies between genus-level reference taxonomies and molecular phylogeny (e.g. *Clostridium* and *Eubacteriumf*)^9^.

In this work, we demonstrate that a substantial improvement in classification accuracy of marker-gene sequences can be achieved if a reference taxonomic distribution for the sample’s source environment is known. This technique enables marker-gene sequencing to differentiate individual species at a level of accuracy previously only available at genus level.

We focus on q2-feature-class¡f¡er, a QIIME 2^10^ plugin for taxonomic classification. In previous work^4^ we benchmarked this method against other common classifiers, including RDP Classifier^11^ and several consensus-based methods using real and simulated data for four bacterial and fungal loci. In general, q2-feature-classifier met or exceeded the accuracy of the other classifiers 4. However, all tested methods performed similarly if their parameters were tuned in a concordant manner. Significant performance enhancement demonstrated in the current work for q2-feature-classifier therefore implies improved performance over those other methods.

RDP Classifier and q2-feature-classifier use similar naive Bayes machine-learning classifiers to assign taxonomies based on sequence k-mer frequencies, and exhibit very similar performance when default parameters are used^4^. The default assumption of these classifiers is that each species in the reference taxonomy is equally likely to be observed. Unlike RDP classifier, however, q2-feature-classifier now allows prior probabilities to be set for each species. We refer to the prior probabilities as *taxonomic weights* and the default equal probabilities as *uniform* weights. We hypothesized that inputting the frequencies with which each taxon is actually observed in nature as taxonomic weights would improve classifier performance.

Taxonomic weights were downloaded and assembled using our new utility, q2-clawback (https://github.com/BenKaehler/q2-clawback). We created weights for 14 Earth Microbiome Project Ontology (EMPO) 3 habitat types^1^ across 21,513 samples from the Qiita microbial study management platform^3^ (see Online Methods for details), q2-clawback can assemble weights from any appropriately curated set of samples or by querying Qiita on any available metadata category. We refer to EMPO 3 habitat-specific taxonomic weights as *bespoke* weights.

Using bespoke weights, researchers can now classify sequences at species level with the same confidence that they previously classified sequences at genus level (Figure 1). The mean error rate (the proportion of reads incorrectly classified) across the 14 EMPO 3 habitat types was 14% (±1% s.e.) for bespoke weights at species level and 16% (±1% s.e.) for uniform weights at genus level (single-sided paired t-test P = 0.14). These results indicate that bespoke weights achieve comparable or better species-level accuracy to what uniform weights can only accomplish at genus level. (As described below, bespoke weights significantly outperform uniform weights by all metrics when both are compared at species level.) The mean Bray-Curtis dissimilarity between observed and expected taxonomic abundances was 0.13 (±0.01 s.e.) for bespoke weights at species level and 0.15 (±0.01 s.e.) for uniform weights at genus level (single-sided paired t-test P = 0.013) (Table S2, Figure S2), indicating better performance of bespoke weights. See Supplementary Results for more details of our benchmarking results, and Online Methods for details.

**Figure 1.**
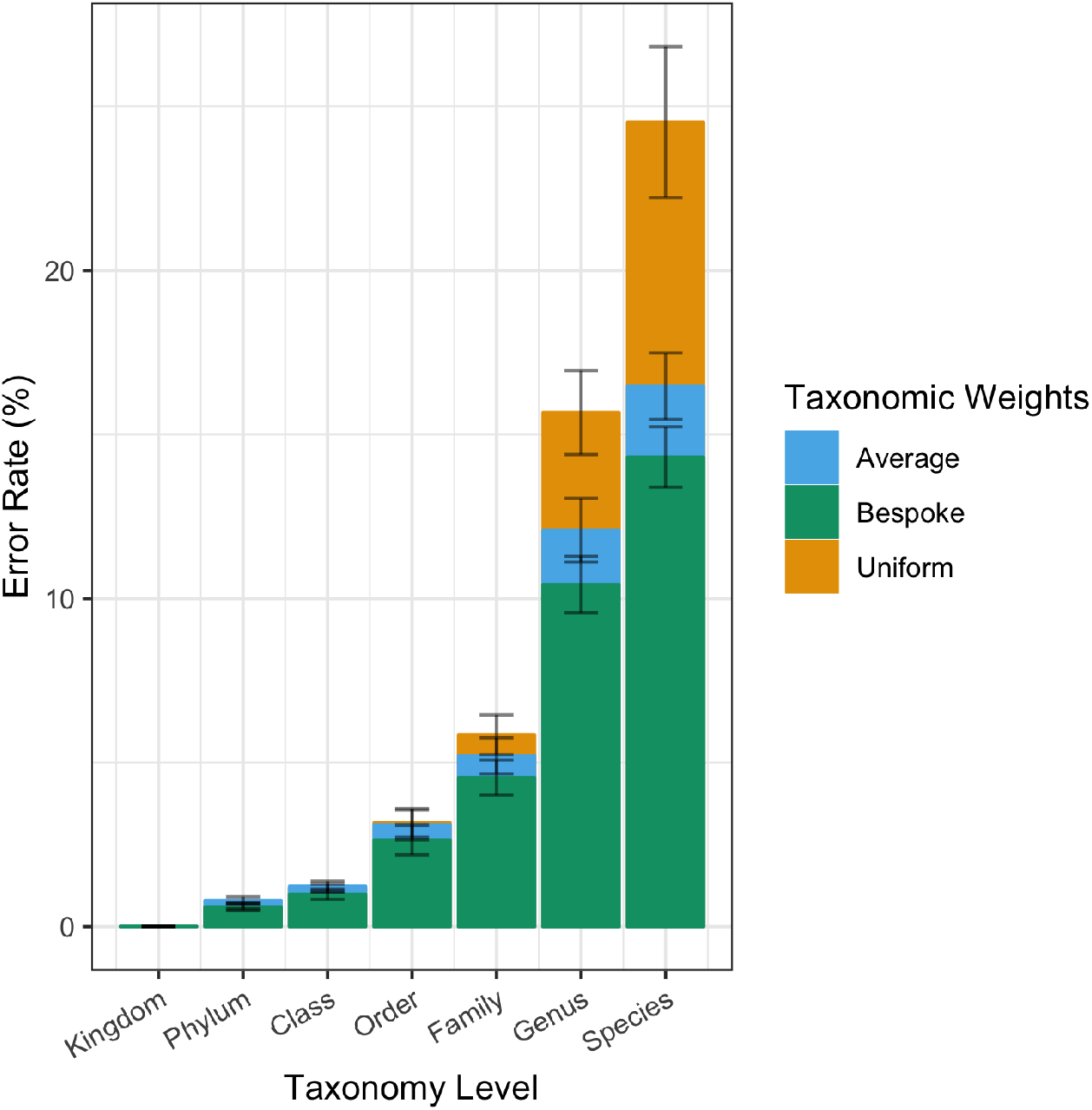
Using habitat-specific taxonomic weights, researchers can now classify sequences at species level with the same confidence that they previously classified sequences at genus level (single-sided paired t-test, species bespoke vs genus uniform, t = 1.6, P = 0.14). Overlaid columns show average proportion of incorrectly classified reads for three taxonomic weighting strategies at each taxonomic level. Bespoke weights were habitat-specific. Average weights were averaged across the 14 EMPO 3 habitats. Uniform weights are the current best practice. Cross validation tests for each of 14 EMPO 3 habitats were averaged to calculated error rates. 21,513 empirical taxonomic abundances contributed to the results. Error bars show standard errors across EMPO 3 habitats.

Bespoke weights significantly outperformed uniform weights when both were compared at species level (bespoke error rate = 14%, uniform error rate = 25%, paired t-test P = 5.8×10^-5^) (Figure 1). Similar results were obtained for Bray-Curtis dissimilarity and F-measure (see Supplementary Results). Averaged across the 14 EMPO 3 habitats, Proteobacteria and Firmicutes were the most abundant phyla (34% and 18% of reads, respectively). Switching from uniform to bespoke weights caused error rates for classification of species in these phyla to drop from 35.4% (±0.7% s.e.) to 22.3% (±0.4% s.e.) and 43.6% (±0.7% s.e.) to 24.3% (±0.3% s.e.) respectively (Figure S8). These differences were highly significant for both Proteobacteria and Firmicutes (paired t-tests P = 1.4×10^-6^ and P= 8.4×10^-6^ respectively).

Classifier performance was sensitive to the choice of taxonomic weights. Testing the use of taxonomic weights from EMPO 3 habitats that were not the sample’s true habitat revealed that as the taxonomic weights moved away from the bespoke weights for a given sample, error rate increased, as expected (Pearson r^2^ = 0.57, P < 2.2×10^-16^) (Figure S5; see Supplementary Results and Online Methods). We also tested the classification accuracy when using the average of the 14 EMPO 3 habitat-specific bespoke weights, which we term *average* weights. For every EMPO 3 habitat, bespoke weights outperformed average weights (sign test P = 6.1×10^−5^) (Figures 2, S2–3). Similarly, average weights always outperformed uniform weights (sign test P = 6.1×10^-5^) (Figures 2, S2–3). The implication is that classification accuracy improves when taxonomic weights more closely resemble taxonomic frequencies observed in nature. Importantly, uniform weights gave inferior performance, even compared to using taxonomic weights from the EMPO 3 habitats other than the sample’s source habitat (*cross-habitat* weights; Figure S4; see Supplementary Results).

**Figure 2.**
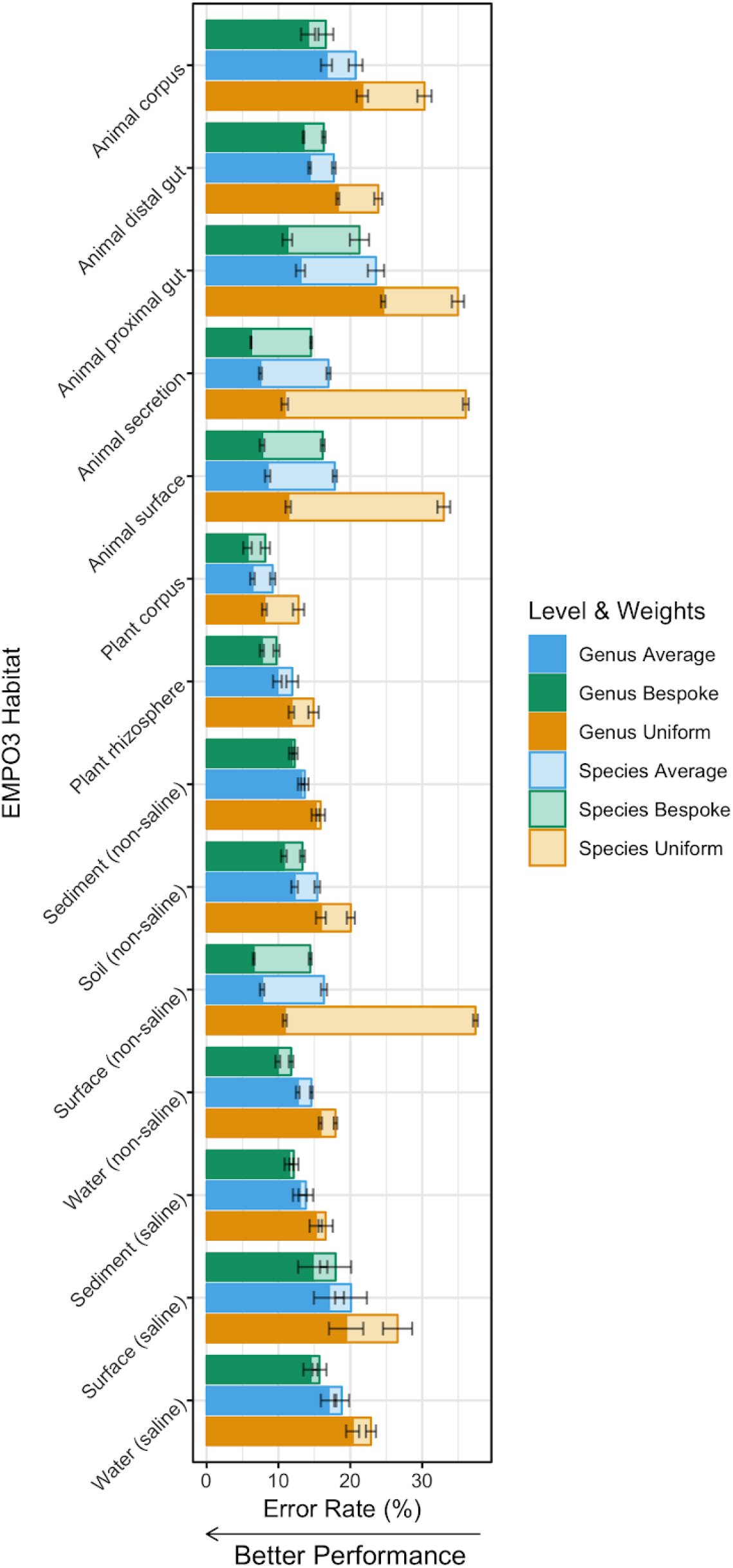
Bespoke weights outperform average weights across EMPO 3 habitat types, and average weights outperform uniform weights (sign test P = 6.1×10^-5^). Columns show average proportion of incorrectly classified reads for differing taxonomic weighting strategies and at genus and species levels. Bespoke weights were habitat-specific. Average weights were averaged across the 14 EMPO 3 habitats. Uniform weights are the current best practice. Tests were based on 5-fold cross validation across 18,222 empirical taxonomic abundances. Error bars show standard errors across folds.

The ability of uniform-weight classifiers to resolve species-level differences from marker genes is directly related to the sequence topology of reference species. Species with highly similar sequences will be difficult to differentiate, even if these species occupy exclusive ecological habitats. However, bespoke weights incorporate habitat-specific species distribution information to guide sequence classification. Hence, classification accuracy under bespoke weights for a given habitat type is tied to the sequence topology and distribution of individual species in that habitat. We devised a statistic that we term the *confusion index* to quantify how often similar sequences originated from different species in the same habitat (see Online Methods). The confusion index is a function of the taxonomic difference between sequences with similar k-mer profiles and the frequency that they appear, taking the bespoke weights as the likelihood of observing a given species. We found that error rates for bespoke weights were correlated with the confusion index (Figure S7; Pearson r^2^ = 0.72, P = 1.4×10^-4^, see Online Methods and Supplementary Results). That is, classification accuracy is affected by how often different species in the same sample have similar amplicon sequences but different taxonomic classifications, and that varies between EMPO 3 habitats.

The assumption of uniform weights, that species are evenly distributed in nature and hence equally likely to be detected, is incorrect. We have demonstrated that this assumption imposes a consistently negative impact on performance, even when compared to deliberately incorrect taxonomic weights selected from ecologically dissimilar environmental sources (the cross-habitat weights). As a result, we suggest the continued usage of uniform weights is not justifiable. When publicly accessible pre-existing microbiome data is available for the sample (i.e., environment) type being investigated, bespoke weights should be used. For all other natural sample types, *average* weights estimated from global microbial species distributions are superior to uniform weights. For highly unusual sample distributions, e.g., in synthetic populations, we recommend compiling custom bespoke weights from existing samples. In the Supplementary Results we demonstrate how shotgun metagenome data may be used to improve classification accuracy (Figure S9). Efforts to curate microbiome data and the continued contribution of researchers to online microbiome data repositories will refine and extend the ability to apply appropriate bespoke weights for sequence classification in diverse sample types.

By comparing uniform, average, and bespoke weights, we have shown that the more specific the taxonomic weights to a sample’s environment, the better the classification accuracy. q2-clawback facilitates achieving these improvements in accuracy by making it easier for the researcher to assemble weights that are more specific than identifying a sample’s EMPO 3 habitat. For instance, it is trivial to assemble weights for all stool samples with human hosts from Qiita (See the online tutorial, https://library.qiime2.org/plugins/q2-clawback).

The results we present provide a general path for delivering species-level classification accuracy. As such, the work provides a complementary solution to the small number of existing specialist classification databases^12–15^. Moreover, bespoke weight classification permits the detection of unexpected species not encompassed by custom databases.

By improving species-level classification of marker-gene sequences, bespoke weights may support critical functional inferences, e.g., differentiation of pathogenic and non-pathogenic species of the same genus^16–21^. Ongoing improvements in public reference sequence and sample databases will further boost performance, supporting biological insight into global microbiome compositions. Uniform weights should always be avoided, as they distort natural species distributions, leading to imprecise and incorrect taxonomic predictions.

## Methods

Methods, including statements of data availability and any associated accession codes and references, are available in the online version of the paper.

## Acknowledgments

QIIME 2 development was primarily funded by NSF Awards 1565100 to JGC and 1565057 to RK. This work was supported by an NHMRC project grant APP1085372, awarded to GAH, JGC, RK.

## Author Contributions

Conceived, designed, and performed experiments: BDK and NAB. Designed and wrote clawback software: BDK and NAB. Wrote manuscript: BDK, NAB, JGC, GAH. Developed supporting software (redbiom): DM, RK. Provided critical review of manuscript and results: DM, RK, JGC, GAH.

## Online Methods

### Data

We downloaded all public 150 nucleotide 16S v4 samples for 18 EMPO 3 habitat types from Qiita^3^ using q2-clawback. The downloaded data consisted of sequence variant and abundance information. The sequence variants were prepared by the standard Qiita pipeline, including Deblur^22^, prior to download. q2-clawback uses redbiom^23^ (https://github.com/biocore/redbiom) to access Qiita. Data from the following Qiita studies were used: 11113^24^, 11444, 1716, 10369^25^, 990^26^, 2080, 1713, 894, 1289, 1883, 1673, 1288, 10353, 2192^27^, 10323, 678, 1773, 662, 1799, 864, 1481, 1024^28^, 1064, 2182, 10934, 1674, 1795^29^, 10273, 10283^30^, 10422^31^, 804, 10308, 1056^32^, 2382^28^, 1240, 889, 1041, 1717, 1222, 11149, 11669, 807^33^, 10245, 1711, 1721, 910, 1001, 895, 550^34^, 1747^35^, 713^36^, 755, 861, 958^37^, 11161^38^, 11154^39^, 945, 723, 1715, 1714, 10798.

The three EMPO 1 control EMPO 3 habitat types were excluded, as well as Hypersaline (saline), Aerosol (non-saline), and Plant surface, which all had fewer than nine samples in the Qiita database for 150 nt sequence variants. The number of samples downloaded for each EMPO 3 habitat are shown in Table OM1.

**Table OM1.** Sample counts for each EMPO 3 habitat type.

| EMPO 3 Habitat Type | Number of Samples |
| --- | --- |
| Animal corpus | 1,158 |
| Animal distal gut | 5,632 |
| Animal proximal gut | 903 |
| Animal secretion | 974 |
| Animal surface | 1,839 |
| Plant corpus | 543 |
| Plant rhizosphere | 328 |
| Sediment (non-saline) | 188 |
| Surface (non-saline) | 1,383 |
| Soil (non-saline) | 2,802 |
| Water (non-saline) | 4,769 |
| Sediment (saline) | 414 |
| Surface (saline) | 152 |
| Water (saline) | 428 |

For the cross validation analysis, sequence-variant level data was discarded and only taxonomic abundance information was retained. The sequence variants were classified using the standard q2-feature-classifier naive Bayes classifier based on Greengenes 99% identity OTU reference data^40^ to obtain empirical taxonomic abundance data for each sample. The naive Bayes classifier was trained using the “balanced” parameter recommendations given in Bokulich, Kaehler, et al.^4^.

For the shotgun data experiment (see Supplementary Results), data was downloaded from the Human Microbiome Project website^2^. The downloaded tables had been prepared using a pipeline leading to MetaPhlAn2^41^. Paired 16S stool samples were downloaded from Qiita^3^ in the form of DNA sequencing data with quality scores. The 16S samples were trimmed to 340 nt and denoised using DADA2^42^. In total, 71 pairs of shotgun and 16S stool samples were found. Reference data sets were downloaded from the NCBI RefSeq database^43^. Full 16S sequences were trimmed to the V3-V5 regions (forward primer CCTACGGGAGGCAGCAG; reverse primer CCGTCAATTCMTTTRAGT), using q2-feature-classifier^4^, resulting in 20,696 reference sequences across 14,777 taxa. It should be noted that this experiment is intended for demonstration only, and that we are not advocating the use of the NCBI 16S RefSeq database for this purpose, as on average there are less than two reference sequence examples for each taxon.

### Clawback

q2-clawback is a free, open-source, BSD-licensed package that is available on GitHub (https://github.com/BenKaehler/q2-clawback). It includes methods for downloading sequence variants from Qiita (fetch-Qiita-samples), extracting sequence variants for taxonomic classification using q2-feature-classifier (sequence-variants-from-samples), and assembling taxonomic weights from collections of samples of taxonomic abundance (generate-class-weights). These methods can be run independently or combined into a single method call (assemble-weights-from-Qiita). Figure OM1 shows the workflow for these methods. An online tutorial is available (https://library.qiime2.org/plugins/q2-clawback.

**Figure OM1.**
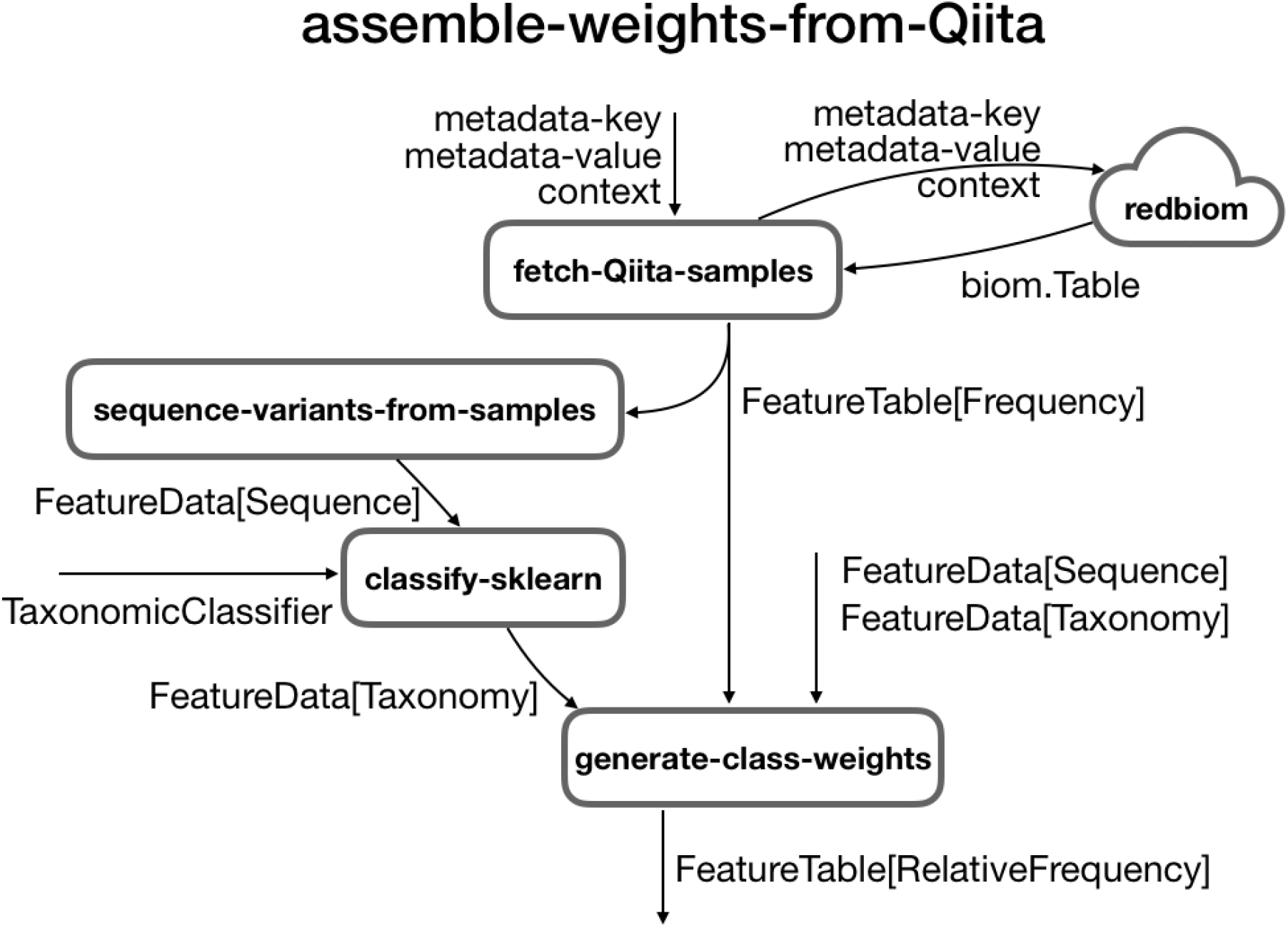
Relationship between q2-clawback methods. q2-clawback contains methods for downloading and assembling taxonomic weights. The assemble-weights-from-Qiita method wraps the illustrated workflow, but fetch-Qiita-samples, sequence-variants-from-samples, and generate-class-weights can also be accessed directly. Labelled data flows show QIIME 2 semantic types and parameters. classify-sklearn is provided by the q2-feature-classifier plugin. redbiom is a service for downloading data from Qiita.

In general, taxonomic weights are assembled as follows. A set of sequence variants with abundances are acquired (fetch-Qiita-samples). The sequence variants are extracted (sequence-variants-from-samples) and classified using the naive Bayes classifier under uniform weights using “balanced” settings^4^. Classification to species level is forced by setting the confidence parameter to −1. The resulting read counts are aggregated, normalised, and added to a small (10^-6^ unobserved weight default) uniform offset (generate-class-weights) to form bespoke weights. The resulting weights are used to retrain the naive Bayes classifier to create a classifier under the bespoke weights assumption. In our experiments, which are detailed below, this procedure was modified slightly to accommodate cross validation and compilation of taxonomic weights from a variety of sources.

### Cross Validation Using Empirical Taxonomic Abundance

To test classification accuracy using varying taxonomic weights, we developed a cross-validation strategy that accounted for the observed abundances of taxa in any given habitat. This strategy ensured that a classifier was never asked to classify a sequence that had occured in its training set or generate taxonomic abundances that had directly contributed to its input taxonomic weights. To our knowledge, our cross-validation strategy is the first to incorporate information about taxonomic weights in assessing taxonomic classifier performance. This situation is known in machine learning as imbalanced learning^44^.

Cross validation was used to analyse the effectiveness of setting the taxonomic weights for the q2-feature-classifier naive Bayes taxonomic classifier. A single cross-validation test follows the pattern (shown in Figure OM2, several steps are described in more detail below):

1. Obtain a set of reference sequences and reference taxonomies.
2. Obtain a set of samples for a given EMPO 3 habitat type, where each sample contains the number of reads observed for each taxon.
3. Perform stratified k-fold cross validation simultaneously on reference sequences and samples.
4. For each fold:

a. Train a classifier on the training reference sequences, optionally incorporating read counts from the training samples to calculate taxonomic weights.
b. Simulate samples that closely match the taxonomic abundances in the test samples using the test reference sequences, then classify them using the above classifier.

**Figure OM2.**
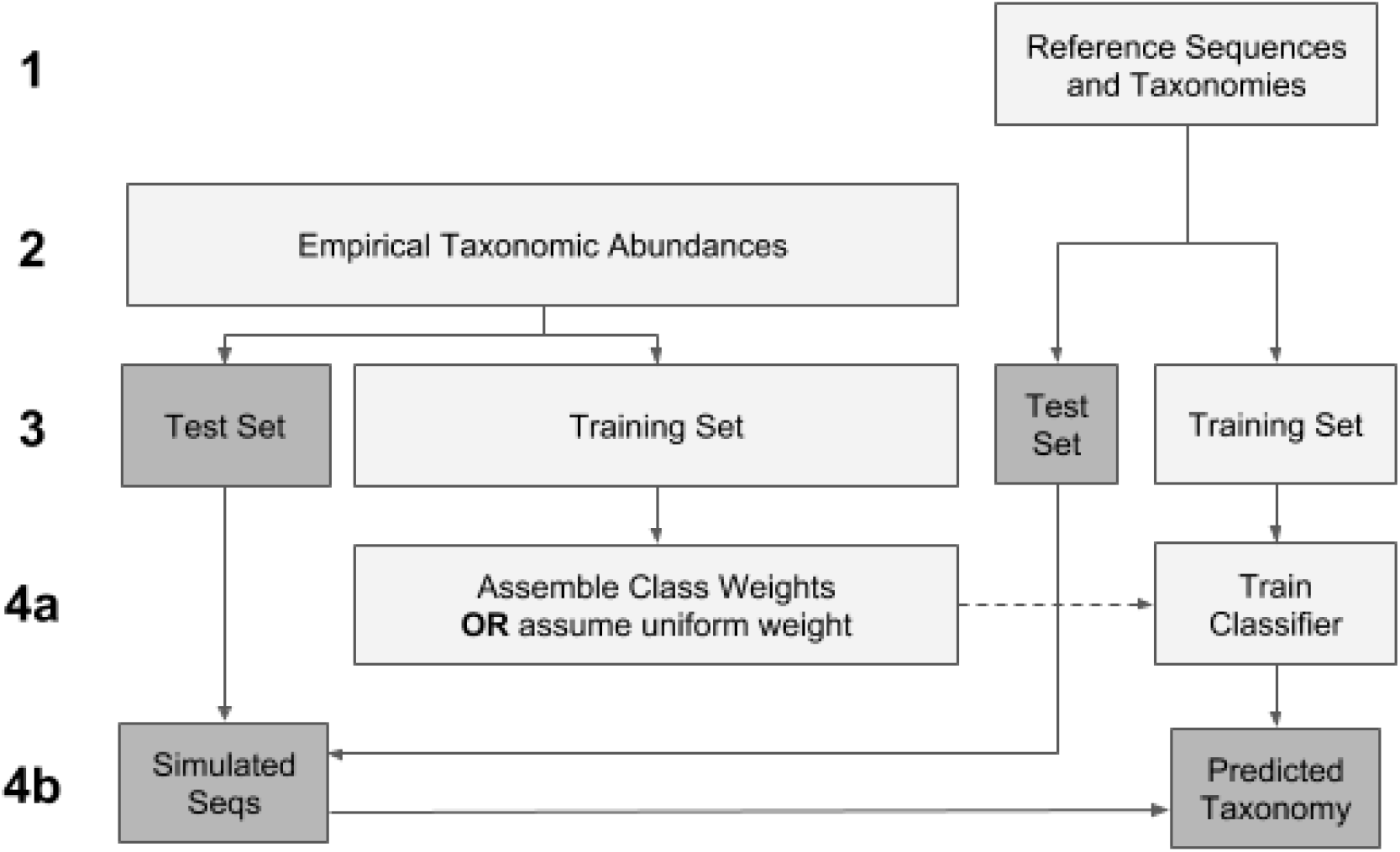
Cross validation workflow. Cross validation on the reference sequences ensured that a classifier was only ever asked to classify unseen sequences. Cross validation on the empirical samples used sequences from the test set of reference sequences to simulate samples with the same taxonomic abundances as the empirical samples, and ensured that bespoke and average weights were never derived from the samples on which they were tested.

**Step 2**. Data was obtained as detailed above. Taxonomic abundances were estimated using the naive Bayes classifier under uniform weights using “balanced” settings^4^, where the classifier was forced to classify to species level.

**Step 3**. We performed 5-fold cross validation in each instance. Standard stratification for 5-fold cross validation requires that at least five sequences exist for each taxonomy, which is not the case for the 99% identity Greengenes reference taxonomy. We therefore formed a stratum for each taxonomy for which five or more reference sequences existed (large taxonomies) and merged the remaining taxonomies (small taxonomies) into those strata. A single large taxonomy was chosen for each small taxonomy by training a naive Bayes classifier on the large taxonomies, classifying the reference sequences in the small taxonomies, then voting weighted by confidence. Shuffled stratified 5-fold cross validation was then implemented using a standard library call to scikit-learn^45^.

Cross validation was performed simultaneously on samples and reference sequences. Sample cross validation was not stratified.

**Step 4a**. Each sample consisted of a set of taxonomies and their abundances. Taxonomic weights were formed by aggregating those counts across the training samples. As a result of the merged strata in Step 3, some taxonomies that were present in the bespoke weights were not present amongst the taxonomies of the training sequences. Any such taxonomy was mapped to the nearest taxonomy that was present amongst the taxonomies represented by the training sequences, as measured by the voting system from Step 3.

**Step 4b**. Samples were simulated by drawing sequences from the test sequences in such a way as to closely resemble the taxonomic abundances of the test samples. Again as a result of the merged strata in Step 3, some taxonomies that were present in the test samples were not present in the taxonomies of the test sequences. In the same way as for Step 4a, any missing species-level taxonomy was mapped to the closest taxonomy for a sequence present in the test sequences. Once missing taxonomies were resolved, samples were simulated by drawing test sequences as evenly as possible from each taxonomy so that any read count was a whole number.

For the q2-feature-classifier naive Bayes classifiers that were reported in this study, we used the recommended “balanced” parameters as recommended for uniform weights^4^. That is, we used a confidence level of 0.7 in all cases. In Bokulich et al.^4^, a confidence level of 0.92 was recommended for bespoke weights tested on mock communities. We tested the classifiers at this level but in all cases the results were dominated by the less conservative confidence level of 0.7.

F-measure and Bray-Curtis^46^ dissimilarity were calculated for each sample and taxonomic level using the q2-quality-control QIIME 2 plugin (https://github.com/qiime2/q2-quality-control). F-measure for each fold was aggregated across samples by weighting by the total read count for each sample. Bray-Curtis dissimilarity was averaged across samples without weighting, but samples with less than 1,000 reads were filtered out.

Error rates, or the proportion of reads not correctly classified, were calculated as follows. A classification was called correct only if the expected classification exactly matched the observed classification to the required taxonomic level. That is, if the expected classification did not contain classification all the way to that level because that species was not present in the training set, then the classification was called correct only if it was truncated at exactly the right level. Correct classification rates were again calculated for each sample and aggregated across samples by weighting by the total read count for each sample. Aggregation across folds and EMPO 3 habitats was evenly weighted.

## Confusion Index

The degree to which species can be successfully resolved is directly related to the dissimilarity of their sequences. We sought to establish a property of the reference data and taxonomic weights that was related to the classification accuracy across EMPO 3 habitats. For any pair of DNA sequences, the critical quantities are their sequence and taxonomic dissimilarities. Sequence dissimilarity is measured as the Bray-Curtis dissimilarity of k-mer counts. Taxonomic dissimilarity is the depth (from species level) of the most recent common ancestor, e.g. zero for the same species, one for species within the same genus and seven for an Archaean versus a Bacterium.

The Confusion Index is then the log of the product of the probability that the sequence dissimilarity for any pair of sequences is less than a threshold (we selected 0.25) and the expectation of the taxonomic distance given that the sequence dissimilarity is less than 0.25. The expectation was calculated under the assumption that the two sequences were sampled independently with probability given by their bespoke weights. That is,

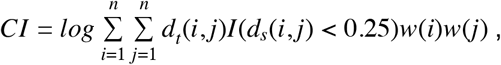

where *CI* is the Confusion Index, *d_s_(i,j)* is the sequence dissimilarity between the *i*th and *j*th sequences, *d_t_(i, j)* is the taxonomic dissimilarity between the *i* th and *j* th sequences, *w(i)* is the weight of the *i* th sequence, and *I*(·) is the indicator function.

The Confusion Index quantifies how often a pair of taxa have nearly identical sequences but different taxonomies for a given set of taxonomic weights. One advantage of this quantity is that it can be estimated statistically by taking a random sample of pairs of sequences. In this study we sampled 10^8^ pairs of sequences for each calculation.

## Comparison of Taxonomic Classification for Shotgun and Amplicon Sequencing

The effect of using taxonomic weights derived from taxonomic classification of shotgun sequencing reads was determined using 5-fold cross validation, where each classifier was trained using taxonomic weights aggregated across the samples in the training set, then tested on 16S samples from a test set. TDR^4^ was computed using the q2-quality-control QIIME 2 plugin. TDR is the fraction of taxa that were discovered in the shotgun sequencing sample that were also found in the amplicon sample.

## Code Availability

q2-clawback is available at https://github.com/BenKaehler/q2-clawback/releases/tag/0.0.4. All other code developed for this study is available at https://github.com/BenKaehler/paycheck/releases/tag/0.0.2.

## Data Availability

The Qiita data used in this study have been deposited at https://doi.org/10.5281/zenodo.2548899. The HMP and NCBI data used in this study have been deposited at https://doi.org/10.5281/zenodo.2549777.

## Supplementary Results for: Species abundance information improves sequence taxonomy classification accuracy

### Cross validation experiments

Using cross validation (see Online Methods), we determined the effect of several different options for obtaining taxonomic weights on taxonomic classification accuracy. We labelled those options as:

– *Uniform weights:* every taxonomic class is assumed to be equally likely.
– *Bespoke weights:* weights drawn from the same EMPO 3 habitat as the test samples.
– *Cross-habitat weights:* weights from any of the 13 EMPO 3 habitats other than a test sample’s source EMPO 3 habitat.
– *Average weights:* weights obtained by averaging across all 14 EMPO 3 habitats.

Uniform weights is the current default assumption for q2-feature-classifier and the only available option for the RDP Classifier. Average weights were used to determine how important it is to closely match the taxonomic weights with the expected weights for a given sample, and to investigate a classification approach for uncharacterized and unknown sample types. Cross-habitat weights were used to determine the effect of classifying reads using misspecified weights. For example, if one were to take a sample that would properly be labelled as Animal distal gut, but erroneously undertake taxonomic classification using a classifier trained using Plant corpus weights.

We used three measures of classification accuracy: error rate, Bray-Curtis dissimilarity, and F-measure, made possible because for each test sample there is a known taxonomy for each read. Please refer to the Online Methods for details of how these measures were calculated and averaged across samples.

### Classification accuracy for bespoke weights at species level meets or exceeds accuracy for uniform weights at genus level

As is typical for existing taxonomy classification methods, classification accuracy was excellent at class level, but decreased at finer levels of taxonomic resolution (Figure S1). Classification accuracy decreased for both bespoke and uniform weighted classifiers, but bespoke classifiers were much less prone to this effect (Figure S1). For uniform weights, the mean F-measure across 14 EMPO 3 habitat types was 0.992 (0.001 standard error), 0.88 (0.01 standard error), 0.81 (0.02 standard error) at class, genus, and species levels, respectively. For bespoke weights, the mean F-measure was 0.992 (0.001 standard error), 0.92 (0.01 standard error), 0.89 (0.01 standard error) at class, genus, and species levels, respectively.

A direct comparison between classification accuracy using bespoke weights versus uniform weights across the 14 EMPO 3 habitat types is given in Tables S1–3, Figure 2, and Figures S2–3 for error rate, F-measure, and Bray-Curtis dissimilarity. In all cases, classification accuracy was better for bespoke weights at species level than for uniform weights at genus level for 10 of the EMPO 3 habitat types (although these 10 habitat types differ among the three measures). The mean error rate (the proportion of reads incorrectly classified) across the 14 EMPO 3 habitat types was 14% (1% standard error) for bespoke weights at species level and 16% (1% standard error) for uniform weights at genus level (single-sided paired t-test P = 0.14) (Figure 1). These results indicate that bespoke weights achieve comparable or better species-level accuracy to what uniform weights can only accomplish at genus level. The mean Bray-Curtis dissimilarity for bespoke weights at species level (0.126, 0.009 standard error) is less than that for uniform weights at genus level (0.15, 0.01 standard error), indicating greater classification accuracy, with a single-sided paired t test P = 0.013. The mean F-measure for bespoke weights at species level (0.887, 0.007 standard error) exceeds that for uniform weights at genus level (0.88, 0.01 standard error) and fails to reject that they are different under a paired t test at 5% significance (P = 0.28). These results verify our claim that on average, classification accuracy at species level for bespoke weights matches or exceeds that for uniform weights at genus level.

### Increasing accuracy of taxonomic weights increases prediction accuracy

Taxonomic classification was tested for uniform, bespoke, average, and cross-habitat taxonomic weights across 14 EMPO 3 habitat types for classification at species level. The results for uniform, bespoke, and average weights are shown for error rate, Bray-Curtis dissimilarity, and F-measure in Figures 2, S2, and S3, respectively. F-measures for cross-habitat taxonomic weights are shown in Figure S4. Cross-habitat results for error rate and Bray-Curtis dissimilarity were similar but are not shown. Note that larger F-measure is better and smaller error rate or Bray-Curtis dissimilarity is better. The total number of samples for each EMPO 3 habitat are shown in Table OM1. The results for bespoke and uniform weights are also summarised in Tables S1–3.

Without exception, for every EMPO 3 habitat and all measures of classification accuracy at genus and species levels, bespoke weights always outperformed average weights, and average weights always outperformed uniform weights (Table S1, Figures 2, S2–3. Across the 14 EMPO 3 habitats, average Bray-Curtis dissimilarities at species level were 0.126 (0.009 standard error), 0.15 (0.01 standard error), 0.23 (0.02 standard error), for bespoke, average, and uniform weights respectively. That is an almost two-fold increase from average bespoke weights Bray-Curtis dissimilarity to that for uniform weights. The average error rates were 14.3% (0.9% standard error), 16% (1% standard error), and 25% (2% standard error), again for bespoke, average, and uniform weights at species level. The corresponding average F-measures were 0.887 (0.007 standard error), 0.871 (0.008 standard error), and 0.81 (0.02 standard error). Note that the variance of the uniform weights results across the EMPO 3 habitats was always greater than for bespoke or average weights. Across the EMPO 3 types, paired t-test differences between bespoke and average weights and average and uniform weights were significant in all cases; maximum P was 4.2×10^-4^. Cross-habitat weights outcomes occupied a spread but rarely outperformed average weights (8 of 182 comparisons); however, cross-habitat weights frequently outperformed uniform weights (117 out of 182 comparisons)(Figure S4). Thus, it appears that uniform weights gravely misrepresent natural species distributions, marring classification accuracy. By comparison, any type of naturally derived taxonomic weight usually improved classification accuracy, even if those weights were derived from a dissimilar habitat type.

Using the cross-habitat weights it was possible to quantify the relationship between classification accuracy and taxonomic weight misspecification over 182 comparisons. We first calculated the differences between error rates using cross-habitat weights and the error rates using bespoke weights. We then calculated the corresponding Kullback-Leibler divergence between the bespoke and cross-habitat weights for each difference and discovered a significant correlation (Pearson r^2^ = 0.57, P < 2.2×10^-16^, see Figure S5). We performed the same test with F-measure and discovered a negative correlation of roughly the same magnitude (Pearson r^2^ = 0.58, P < 2.2×10^-16^). In our tests, using the bespoke weights yielded the best classification accuracy at every level, regardless of how we measured it. This result refines that finding to show that the amount by which performance degrades for taxonomic weights other than the default weights is proportional to how different they are to the bespoke weights. The implication is that any improvement of uniform weights in the direction of bespoke weights is worthwhile, and that if bespoke weights are not available, then taxonomic weights from a similar habitat or the average weights should be used for classification.

### Classification performance is largely predictable based on weights and reference data

We investigated what factors lead the classification accuracy for bespoke weights to vary between EMPO 3 habitats. To give an idea of the shapes of the taxonomic weights distributions, the cumulative distributions of taxonomic weights for each EMPO 3 habitat type are shown in Figure S6. The distributions are qualitatively similar, with the most abundant 500 out of 5,403 species for each habitat accounting for greater than 93% of the mass in each case. While it is not shown in Figure S6, it is also worth noting that the most common taxa for each habitat type are similar: the union of the sets of taxa that account for the first 95% of weights for each habitat type contains only 1,571 taxa.

We first examined whether the diversity of an EMPO 3 habitat was related to classification accuracy under bespoke weights. Regression of error rate against the entropy of the taxonomic weights for each of the 14 EMPO 3 habitats showed no significant relationship (Pearson r^2^ = 0.12, P = 0.23).

The classification accuracy for a given EMPO 3 habitat type was instead found to be largely explained by specific interaction between taxonomic weights and the topology of the space of sequences. We discovered a significant correlation between the confusion index (see Online Methods) and the error rate using bespoke weights for each of the 14 EMPO 3 types (Pearson r^2^ = 0.72, P = 1.3×10^-4^) (Figure S7). A negative correlation of a similar magnitude was found for F-measure (Pearson r^2^ = 0.63, P = 7.1×10^-4^). While much has been written about the difficulty of establishing species-level identity from short read marker-gene sequences^4–8^, to our knowledge this is the first instance of a systematic quantitative analysis of how difficult this problem is in the light of which species co-occur. By using typical weights (as estimated from the data) we have shown that while it is possible to find many examples where a taxonomic classifier can be confused at species level if presented with isolated genetic sequences and the entire reference database, that in practice the problem does not have to be that hard. More than explaining variation between habitat types, these discoveries give a clear indication of the basis for the improvements that we see when using bespoke weights.

### Classification performance varies by phylum

The error rate was tested for each of the phyla present in the observed taxa across all 14 EMPO 3 habitats under the assumptions of uniform and bespoke weights. The results for the most abundant phyla (those with average abundance > 0.5%) are shown in Figure S8, where error rate and abundance was averaged over the habitat types. We observed that reads from some phyla are significantly more difficult to classify than others. For instance, using uniform weights, the error rate was 44% (0.7% standard error) for Firmicutes but 4% (0.2% standard error) for Acidobacteria. For the two most abundant phyla, Proteobacteria and Firmicutes, the decrease in incorrect classifications from uniform to bespoke weights was substantial, from 35% (0.7% standard error) to 22% (0.4% standard error) and from 44% (0.7% standard error) to 24% (0.3% standard error), respectively (maximum t-test P = 8.4×10^-6^). For Firmicutes that is an almost two-fold reduction in the number of incorrectly classified reads. These increases in accuracy underline the consistent increases in accuracy from uniform to bespoke weights that we have observed throughout this study.

### Amplicon sequencing as a proxy for shotgun metagenomics

Shotgun sequencing data, which has the potential to be less biased and higher-resolution than short amplicon sequences^47^, may provide high-accuracy taxonomic weights to further increase the value of high-throughput amplicon sequence data. To test this hypothesis, we downloaded 71 stool samples from the Human Microbiome Project website^2^ for which shotgun and marker-gene data were available. Again using cross validation and treating the shotgun sequencing taxonomic classifications as ground-truth, the taxon discovery rate (TDR)^4^ for species-level classification of denoised 16S rRNA gene sequences improved from 0.46 (0.009 standard error) to 0.54 (0.01 standard error) when using shotgun-derived taxonomic weights relative to uniform weights (paired t-test P = 1.4×10^-20^). TDR using shotgun weights at species level also exceeded that using uniform weights at genus level (0.50, 0.009 standard error) (paired t-test P = 3.9×10^-5^).

Note that this is also a verification of our findings that does not use cross validation on reference sequences or the Greengenes^40^ reference taxonomy.

## Supplementary Tables and Figures

**Table S1.** Using habitat-specific taxonomic weights, researchers can now classify sequences at species level with the same confidence that they previously classified sequences at genus level. Table shows error rates at genus and species levels for habitat-specific (bespoke) and standard (uniform) taxonomic weights. Bolded rows indicate EMPO 3 habitats where species-level error rate with the bespoke classifier is less than genus-level accuracy with the uniform classifier. Lower error rate indicates superior accuracy.

|  | Error Rate (%)* |  |  |  |
| --- | --- | --- | --- | --- |
|  | Bespoke |  | Uniform |  |
|  | Genus | Species | Genus | Species |
| EMPO 3 Habitat |  |  |  |  |
| **Animal corpus** | **14.1** | **16.6** | **21.7** | **30.3** |
| **Animal distal gut** | **13.5** | **16.3** | **18.3** | **23.9** |
| **Animal proximal gut** | **11.2** | **21.3** | **24.6** | **35.0** |
| Animal secretion | 6.2 | 14.5 | 10.9 | 36.1 |
| Animal surface | 7.7 | 16.2 | 11.4 | 33.0 |
| Plant corpus | 5.7 | 8.2 | 8.0 | 12.8 |
| **Plant rhizosphere** | **7.7** | **9.7** | **11.8** | **14.9** |
| **Sediment (non-saline)** | **11.8** | **12.3** | **15.2** | **15.9** |
| **Soil (non-saline)** | **10.8** | **13.3** | **15.9** | **20.1** |
| Surface (non-saline) | 6.5 | 14.4 | 10.9 | 37.4 |
| **Water (non-saline)** | **9.9** | **11.8** | **15.8** | **17.9** |
| **Sediment (saline)** | **11.5** | **12.1** | **15.2** | **16.6** |
| **Surface (saline)** | **14.8** | **17.9** | **19.4** | **26.6** |
| **Water (saline)** | **14.5** | **15.7** | **20.3** | **22.9** |

\*Error rates (proportion of reads incorrectly classified) are from 5-fold cross validation where classifiers were tested on sequences and empirical taxonomic abundances that were not used in training. Tests were based on 21,513 empirical samples across the 14 habitat types.

**Table S2.**
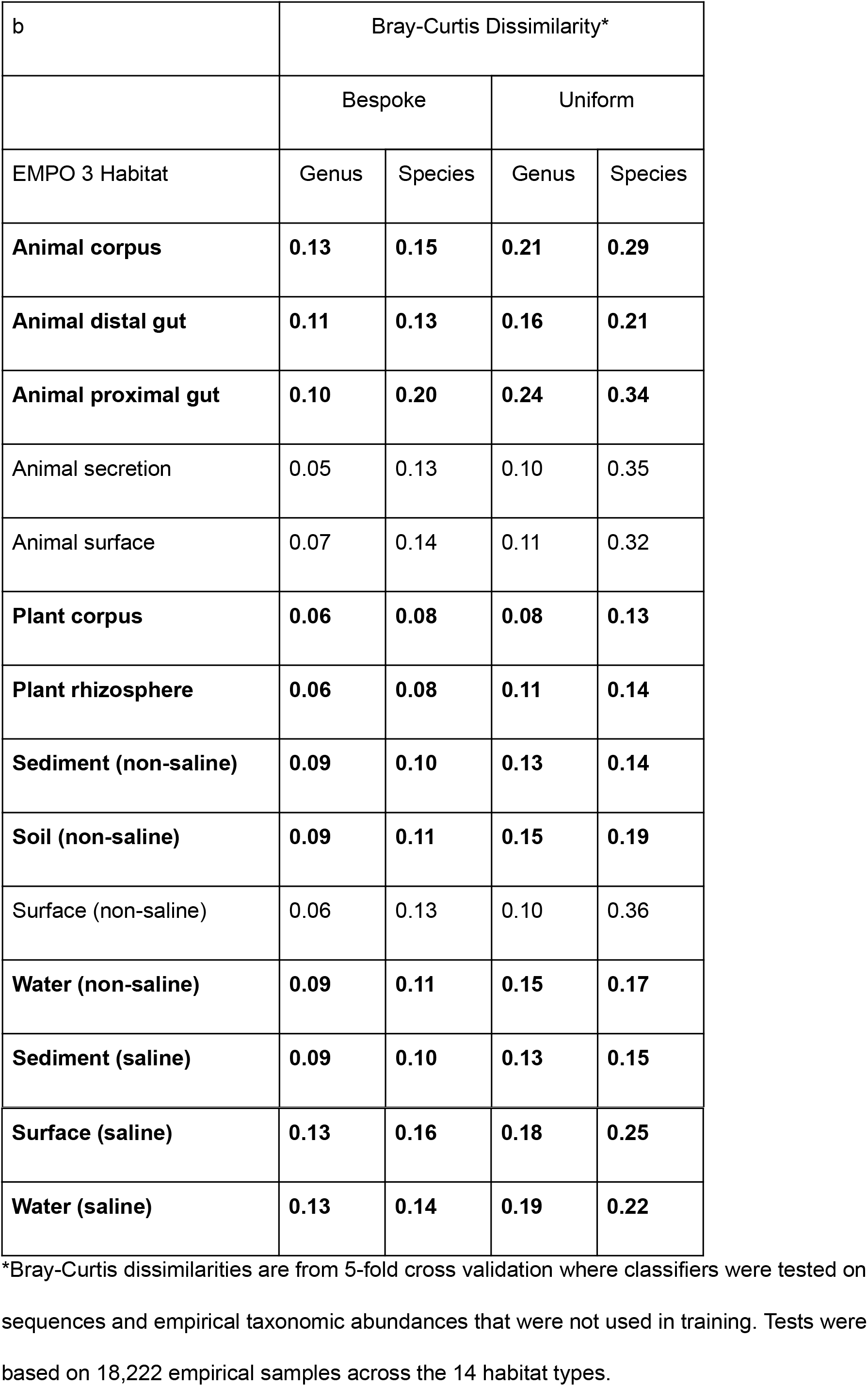
Using habitat-specific taxonomic weights, researchers can now classify sequences at species level with the same confidence that they previously classified sequences at genus level. Table shows Bray-Curtis dissimilarity at genus and species levels for habitat-specific (bespoke) and standard (uniform) taxonomic weights. Bolded rows indicate EMPO 3 habitats where species-level Bray-Curtis dissimilarity with the bespoke classifier is less than genus-level accuracy with the uniform classifier. Lower Bray-Curtis dissimilarity indicates superior accuracy.

| b | Bray-Curtis Dissimilarity* |  |  |  |
| --- | --- | --- | --- | --- |
|  | Bespoke |  | Uniform |  |
| EMPO 3 Habitat | Genus | Species | Genus | Species |
| **Animal corpus** | **0.13** | **0.15** | **0.21** | **0.29** |
| **Animal distal gut** | **0.11** | **0.13** | **0.16** | **0.21** |
| **Animal proximal gut** | **0.10** | **0.20** | **0.24** | **0.34** |
| Animal secretion | 0.05 | 0.13 | 0.10 | 0.35 |
| Animal surface | 0.07 | 0.14 | 0.11 | 0.32 |
| **Plant corpus** | **0.06** | **0.08** | **0.08** | **0.13** |
| **Plant rhizosphere** | **0.06** | **0.08** | **0.11** | **0.14** |
| **Sediment (non-saline)** | **0.09** | **0.10** | **0.13** | **0.14** |
| **Soil (non-saline)** | **0.09** | **0.11** | **0.15** | **0.19** |
| Surface (non-saline) | 0.06 | 0.13 | 0.10 | 0.36 |
| **Water (non-saline)** | **0.09** | **0.11** | **0.15** | **0.17** |
| **Sediment (saline)** | **0.09** | **0.10** | **0.13** | **0.15** |
| **Surface (saline)** | **0.13** | **0.16** | **0.18** | **0.25** |
| **Water (saline)** | **0.13** | **0.14** | **0.19** | **0.22** |

\*Bray-Curtis dissimilarities are from 5-fold cross validation where classifiers were tested on sequences and empirical taxonomic abundances that were not used in training. Tests were based on 18,222 empirical samples across the 14 habitat types.

**Table S3.** Using habitat-specific taxonomic weights, researchers can now classify sequences at species level with the same confidence that they previously classified sequences at genus level. Table shows F-measure at genus and species levels for habitat-specific (bespoke) and standard (uniform) taxonomic weights. Bolded rows indicate EMPO 3 habitats where species-level F-measure with the bespoke classifier is greater than genus-level accuracy with the uniform classifier. Greater F-measure indicates superior accuracy.

| c | F-measure* |  |  |  |
| --- | --- | --- | --- | --- |
|  | Bespoke |  | Uniform |  |
| EMPO 3 Habitat | Genus | Species | Genus | Species |
| **Animal corpus** | **0.89** | **0.87** | **0.83** | **0.76** |
| **Animal distal gut** | **0.89** | **0.87** | **0.85** | **0.81** |
| **Animal proximal gut** | **0.91** | **0.83** | **0.81** | **0.72** |
| Animal secretion | 0.95 | 0.89 | 0.92 | 0.73 |
| Animal surface | 0.94 | 0.88 | 0.92 | 0.76 |
| Plant corpus | 0.95 | 0.93 | 0.94 | 0.90 |
| **Plant rhizosphere** | **0.94** | **0.92** | **0.91** | **0.89** |
| **Sediment (non-saline)** | **0.91** | **0.90** | **0.88** | **0.87** |
| **Soil (non-saline)** | **0.92** | **0.90** | **0.88** | **0.85** |
| Surface (non-saline) | 0.95 | 0.89 | 0.92 | 0.73 |
| **Water (non-saline)** | **0.93** | **0.91** | **0.88** | **0.86** |
| **Sediment (saline)** | **0.91** | **0.90** | **0.88** | **0.87** |
| **Surface (saline)** | **0.89** | **0.85** | **0.85** | **0.79** |
| **Water (saline)** | **0.89** | **0.87** | **0.84** | **0.81** |

\*F-measures are for 5-fold cross validation where classifiers were tested on sequences and empirical taxonomic abundances that were not used in training. Tests were based on 21,513 empirical samples across the 14 habitat types.

**Figure S1.**
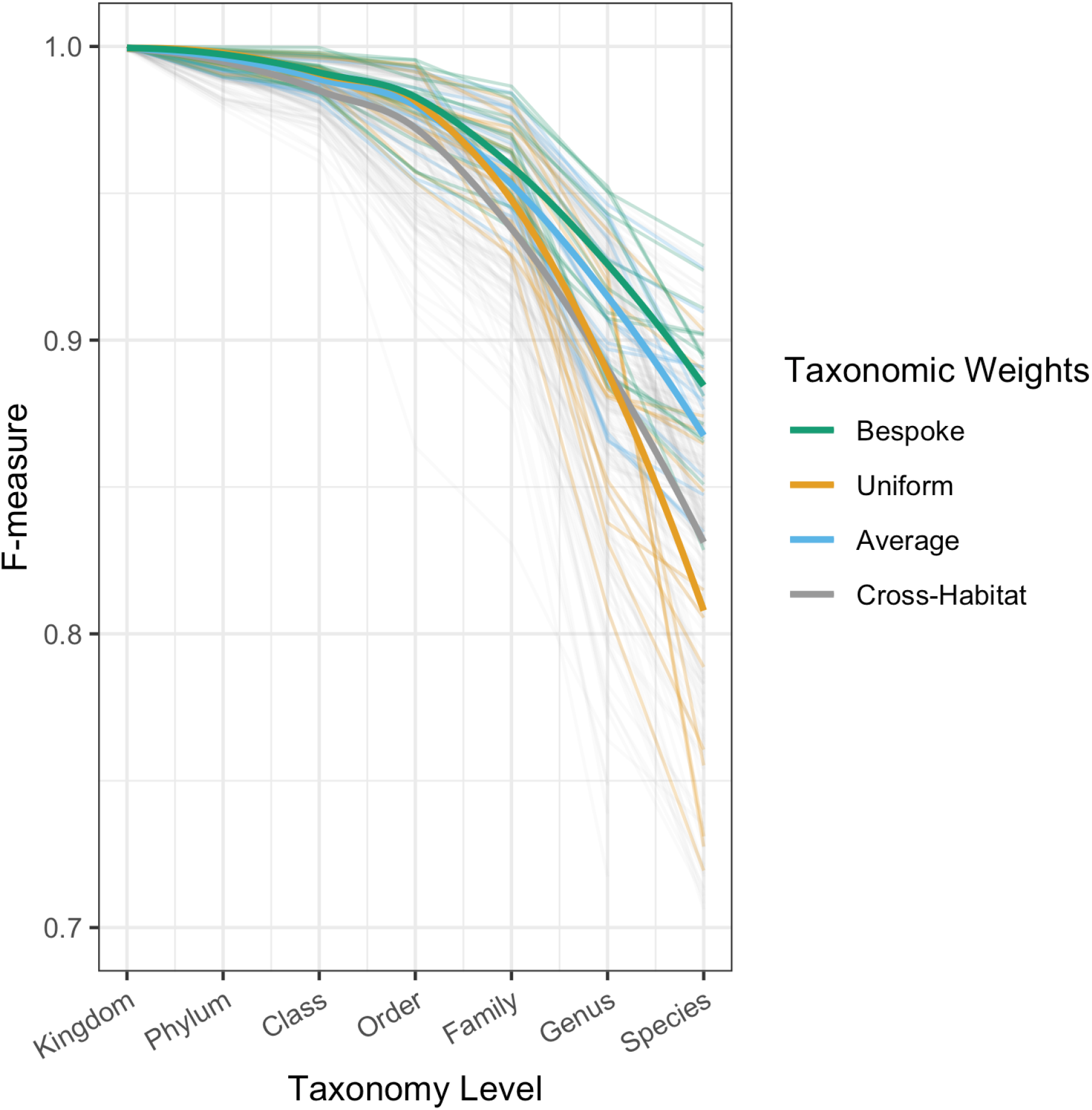
Classification accuracy drops with increasing taxonomic specificity. F-measures are for 5-fold cross validation where classifiers trained using a variety of taxonomic weighting strategies are tested on sequences and empirical taxonomic abundances that were not used in training. Classification F-measure drops as finer levels of classification are required, but is much more consistent across levels for classifiers with bespoke (habitat-specific) weights. Bespoke weights were habitat-specific. Average weights were averaged across the 14 EMPO 3 habitats. Uniform weights are the current best practice. Cross-habitat weights were weights from EMPO 3 habitats other than the sample’s habitat. Faint lines show results for 14 EMPO 3 habitat types. Bold lines show LOESS plots to demonstrate trends.

**Figure S2.**
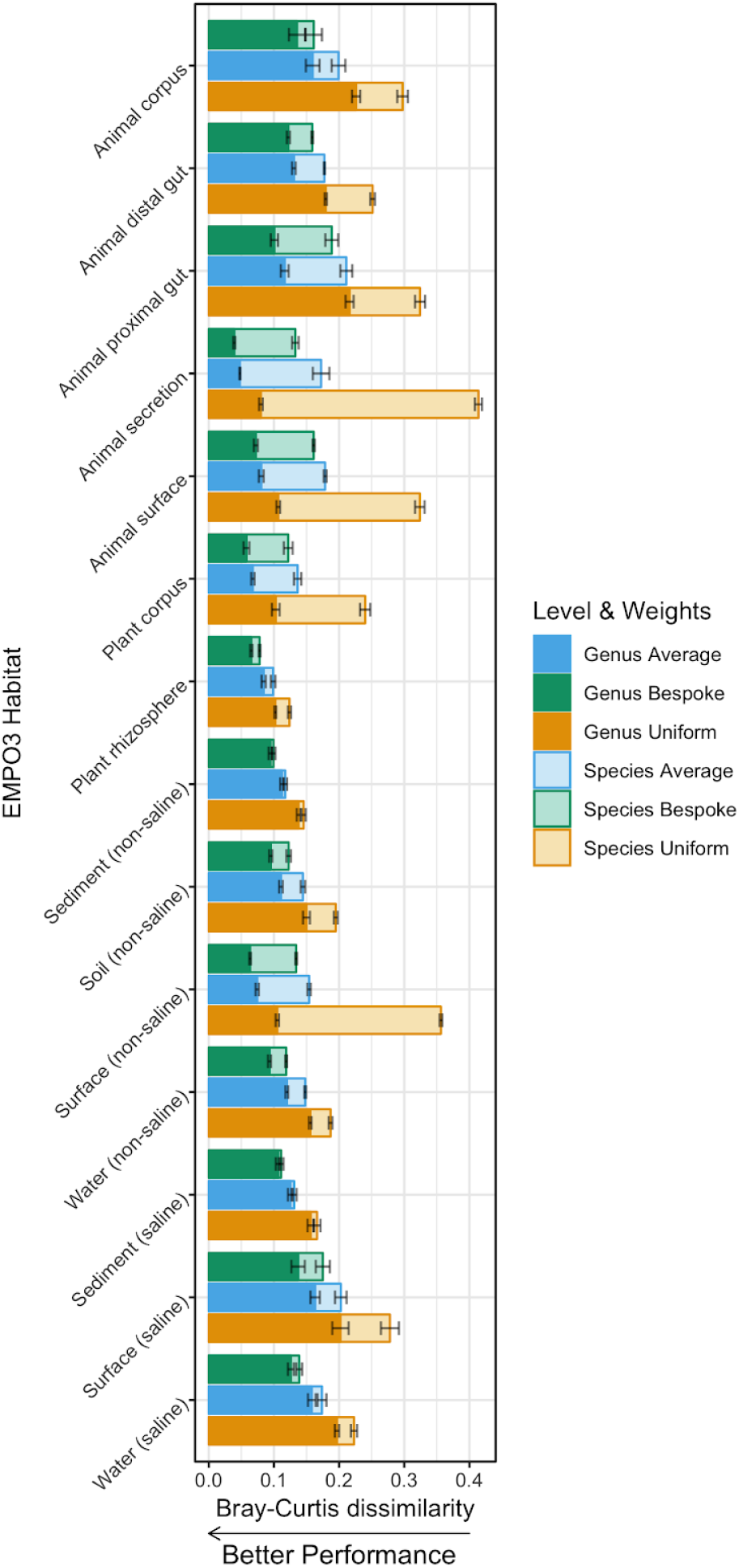
Bespoke weights always outperformed average weights across EMPO 3 habitat types, and average weights always outperformed uniform weights (sign test P = 6.1×10^-5^). Columns show average Bray-Curtis dissimilarity between expected and observed taxonomic abundances for differing taxonomic weighting strategies and at genus and species levels. Bespoke weights were habitat-specific. Average weights were averaged across the 14 EMPO 3 habitats. Uniform weights are the current best practice. Tests were based on 5-fold cross validation across 18,222 empirical taxonomic abundances. Error bars show standard errors across folds.

**Figure S3.**
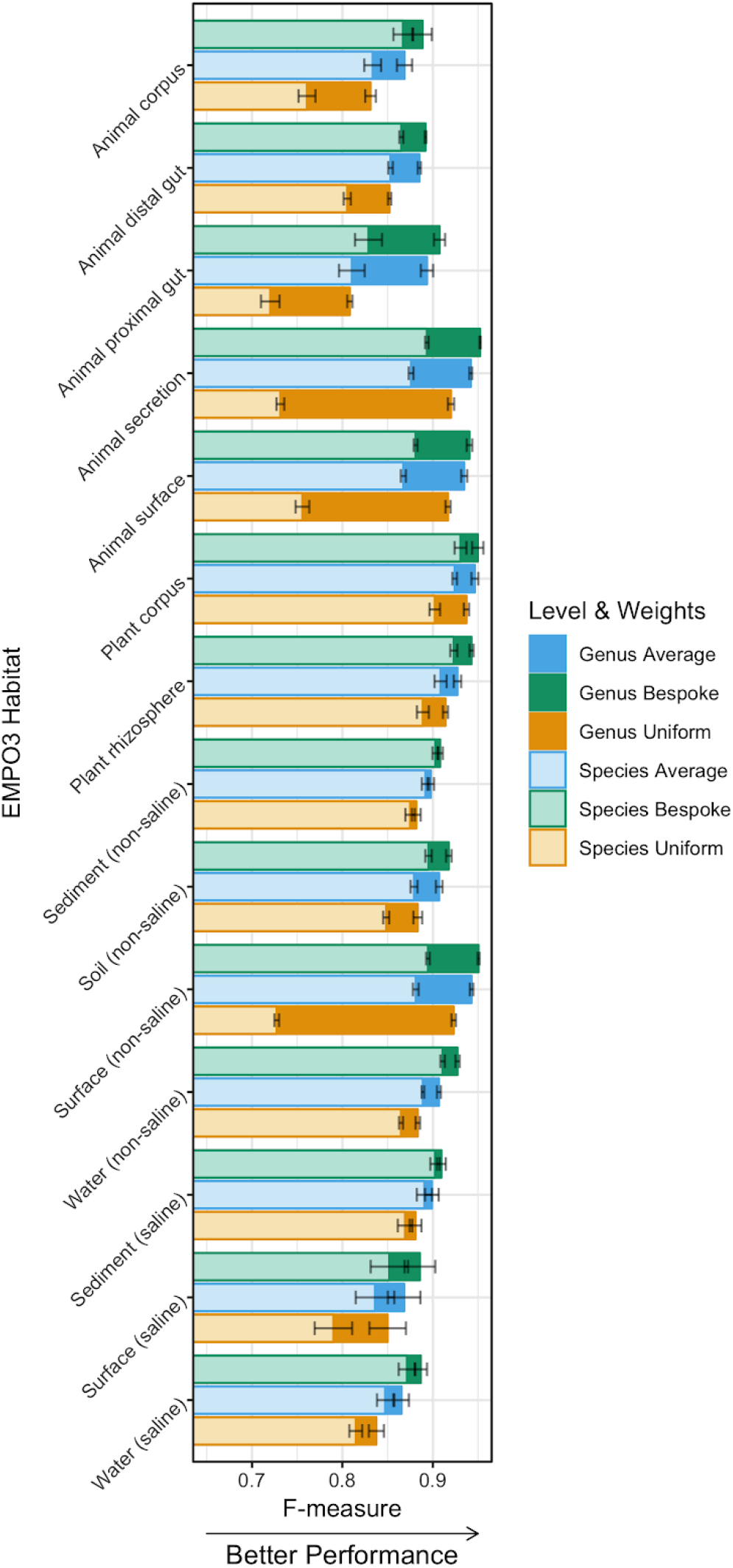
Bespoke weights always outperformed average weights across EMPO 3 habitat types, and average weights always outperformed uniform weights (sign test P = 6.1×10^-5^). F-measures are from 5-fold cross validation where classifiers trained using a variety of taxonomic weighting strategies are tested on sequences and empirical taxonomic abundances that were not used in training. Tests were based on 21,513 empirical samples across the 14 habitat types. Error bars show standard errors across folds.

**Figure S4.**
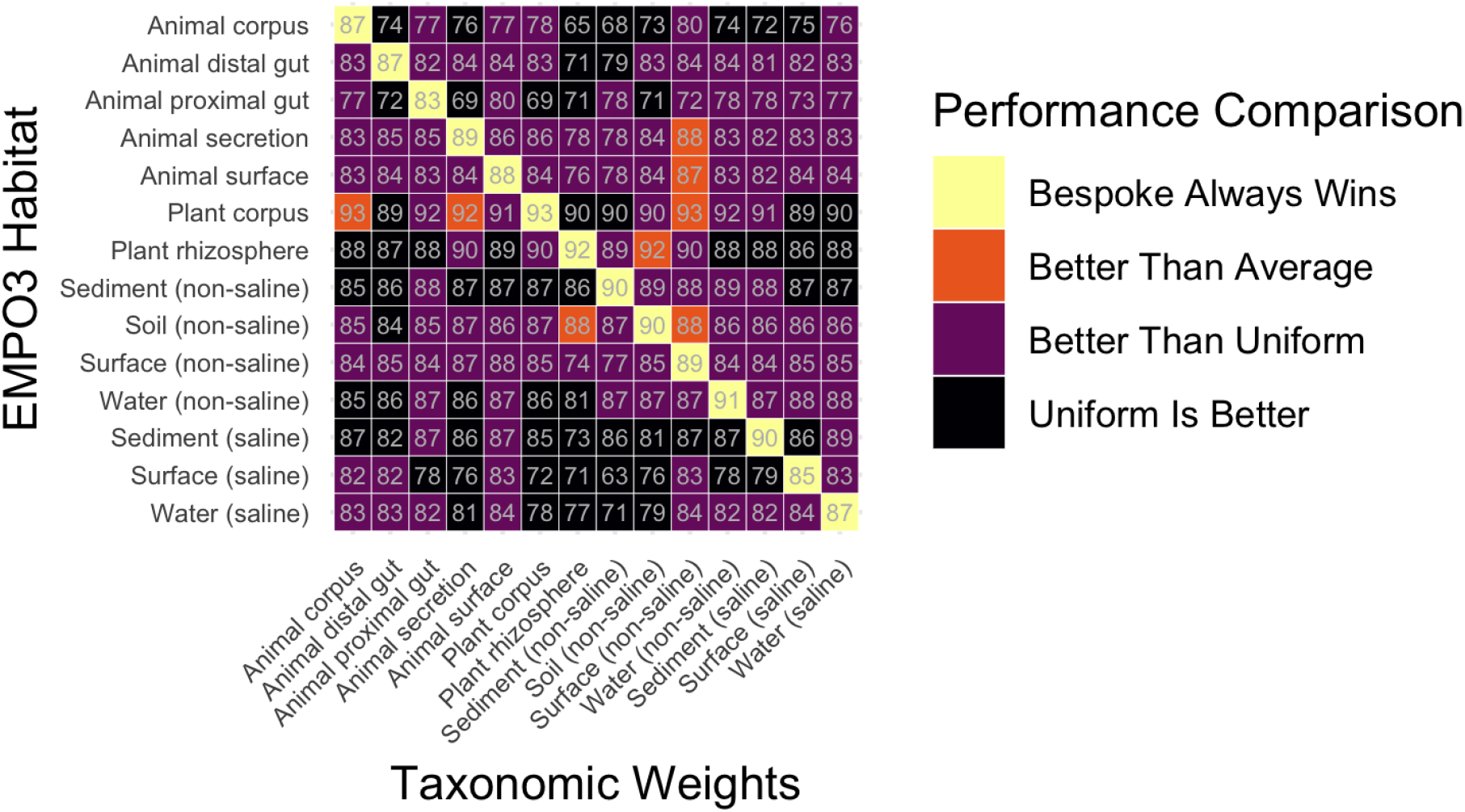
Summary of the effect of using cross-habitat weights (taxonomic weights different to the sample habitat). Light grey numbers show F-measure as a percentage. Bespoke weights (when taxonomic weights match sample habitat) are always superior. Occasionally (8 times) weights other than bespoke weights beat the weights that were averaged across the 14 EMPO 3 habitats. Frequently the cross-habitat weights were better than uniform weights (which is current best practice, 109 times). Occasionally the uniform weights outperformed the cross-habitat weights (65 times).

**Figure S5.**
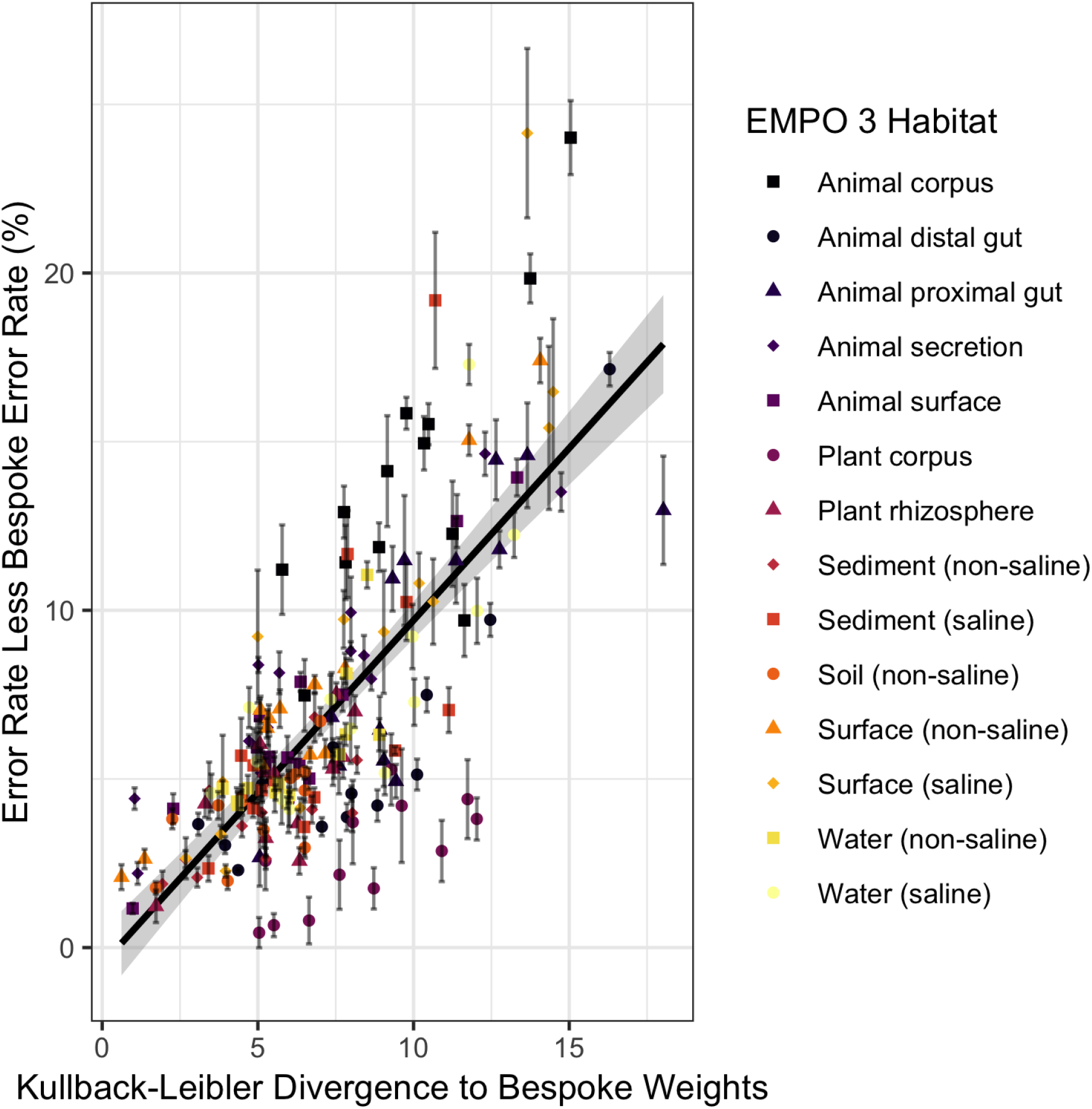
Classification accuracy degrades as taxonomic weights diverge from sample abundances. Cross testing of classification accuracy by setting taxonomic weights to those from each of the 13 EMPO 3 habitats other than the appropriate bespoke weights, across 14 EMPO 3 habitats. There is a clear association (Pearson r^2^ = 0.57, P < 2.2×10^-16^).

**Figure S6.**
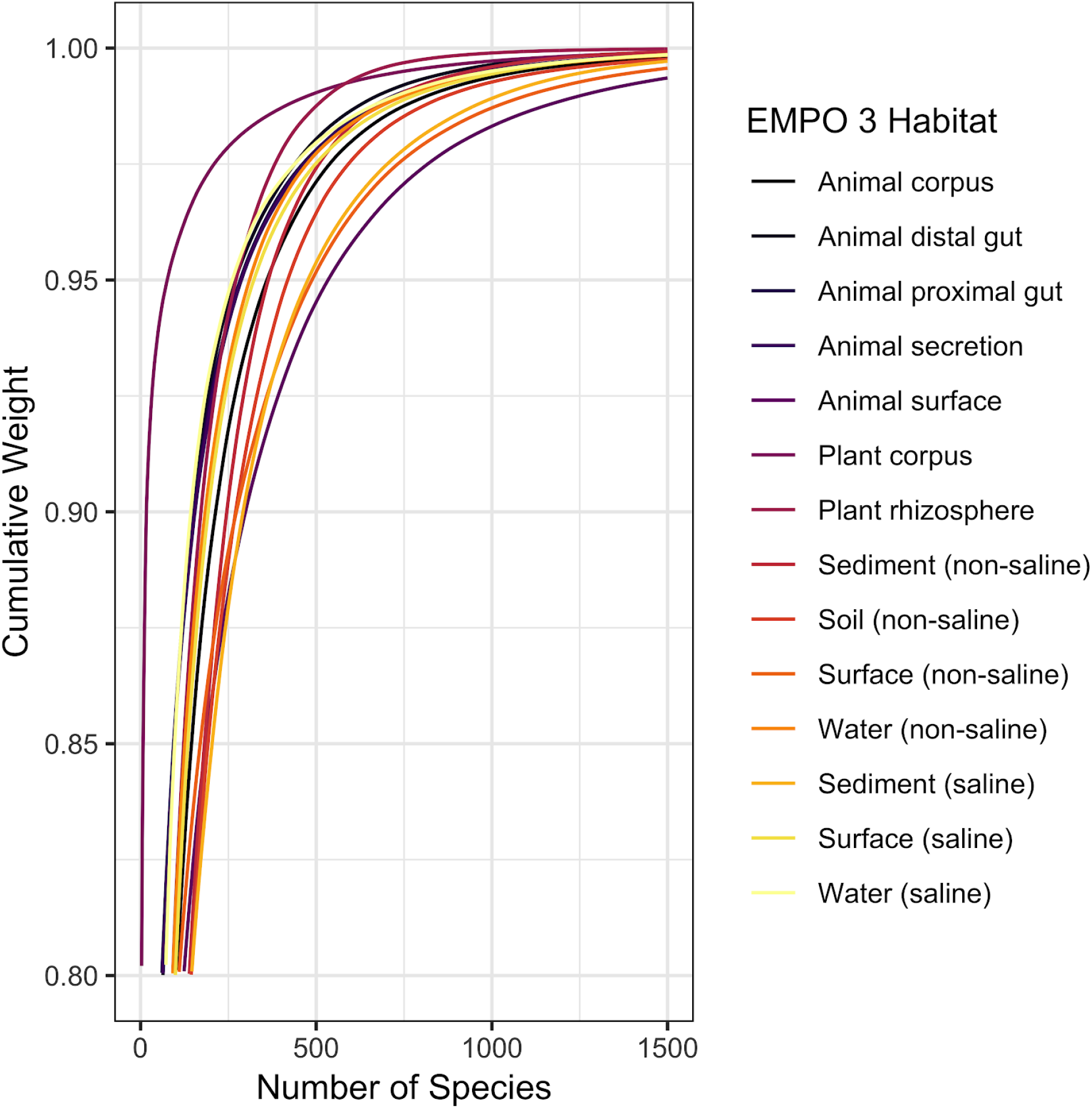
Cumulative weights for different habitat types display similar diversity trends. Lines show the cumulative taxonomic weights for 14 EMPO 3 habitat types as a function of species count, coloured by habitat. Taxa are ordered separately for each habitat from most to least abundant. The most peaked distribution is Plant corpus, where 73% of reads were mapped to a single taxonomy in the Cyanobacteria phylum, Chloroplast class.

**Figure S7.**
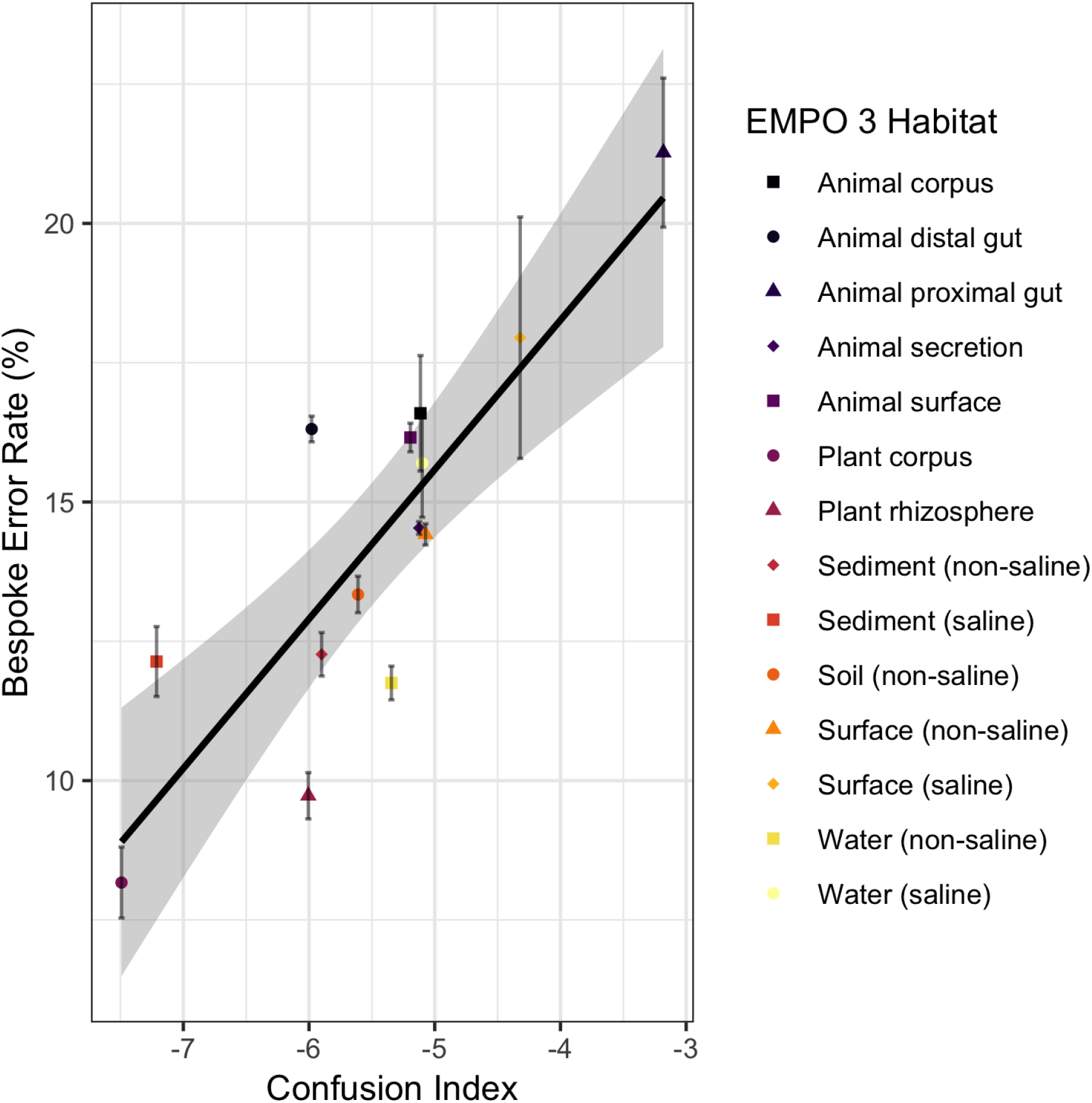
Classification accuracy when using the appropriate bespoke weights is largely explained by how often sequences from different from species are confused (Pearson r^2^ = 0.72, P = 1.3×10^-4^). The confusion index is the log of the expected level of taxonomic difference between two similar reference sequences weighted by the likelihood of observing similar sequences. All points calculated using 5-fold cross validation. Error bars are standard errors across folds. Regression confidence intervals are 95%.

**Figure S8.**
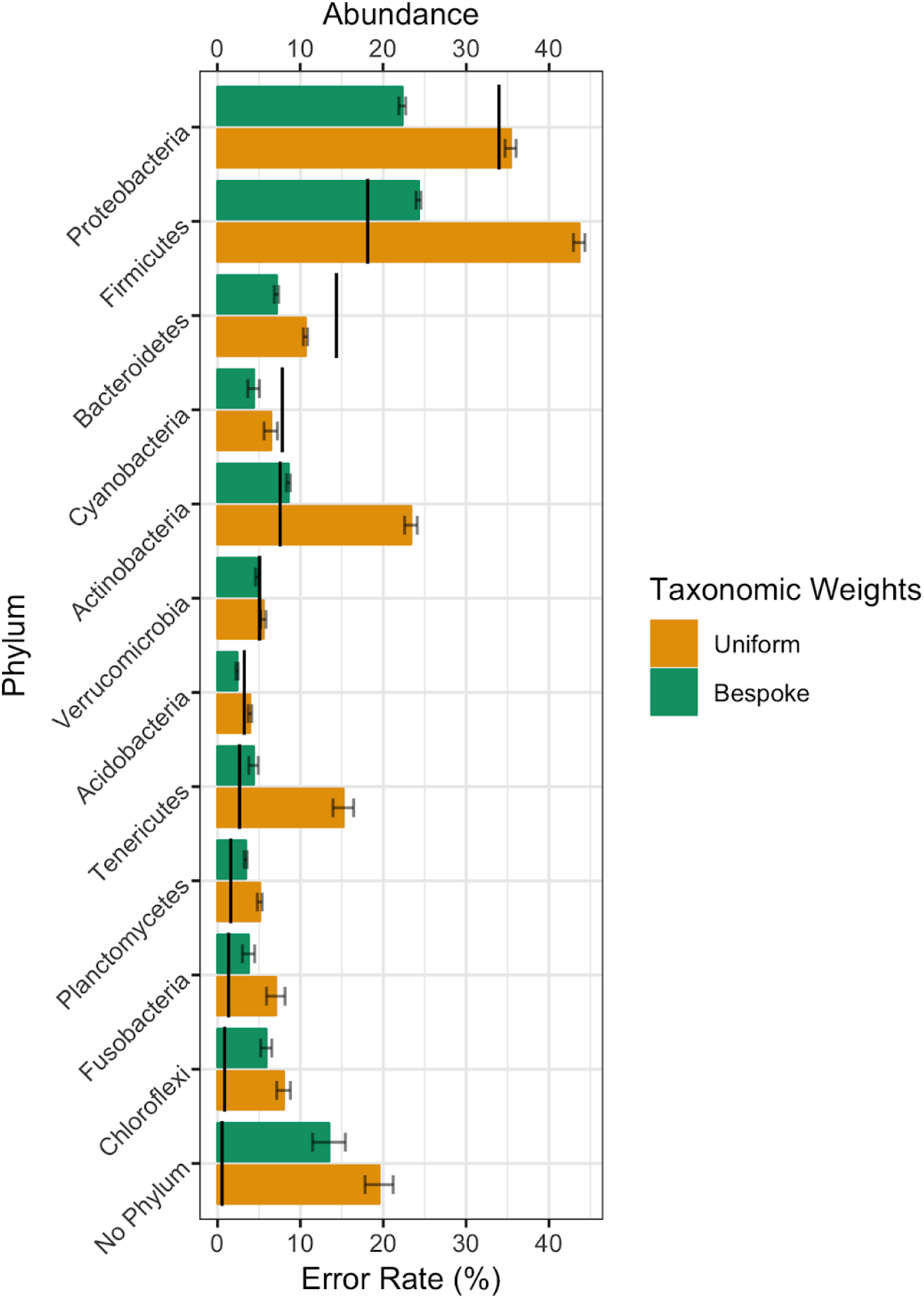
Habitat-specific taxonomic weights improve species-level classification accuracy across phyla. The use of habitat-specific weights is more important for species classification within some phyla, but is more important for more abundant phyla. Columns show percentage of reads correctly classified averaged across 14 EMPO 3 habitats and 21,513 empirical samples. Black lines show average abundances for each phylum. The phyla were truncated to only show those with an average abundance of > 0.05%. Error bars show standard error.

**Figure S9.**
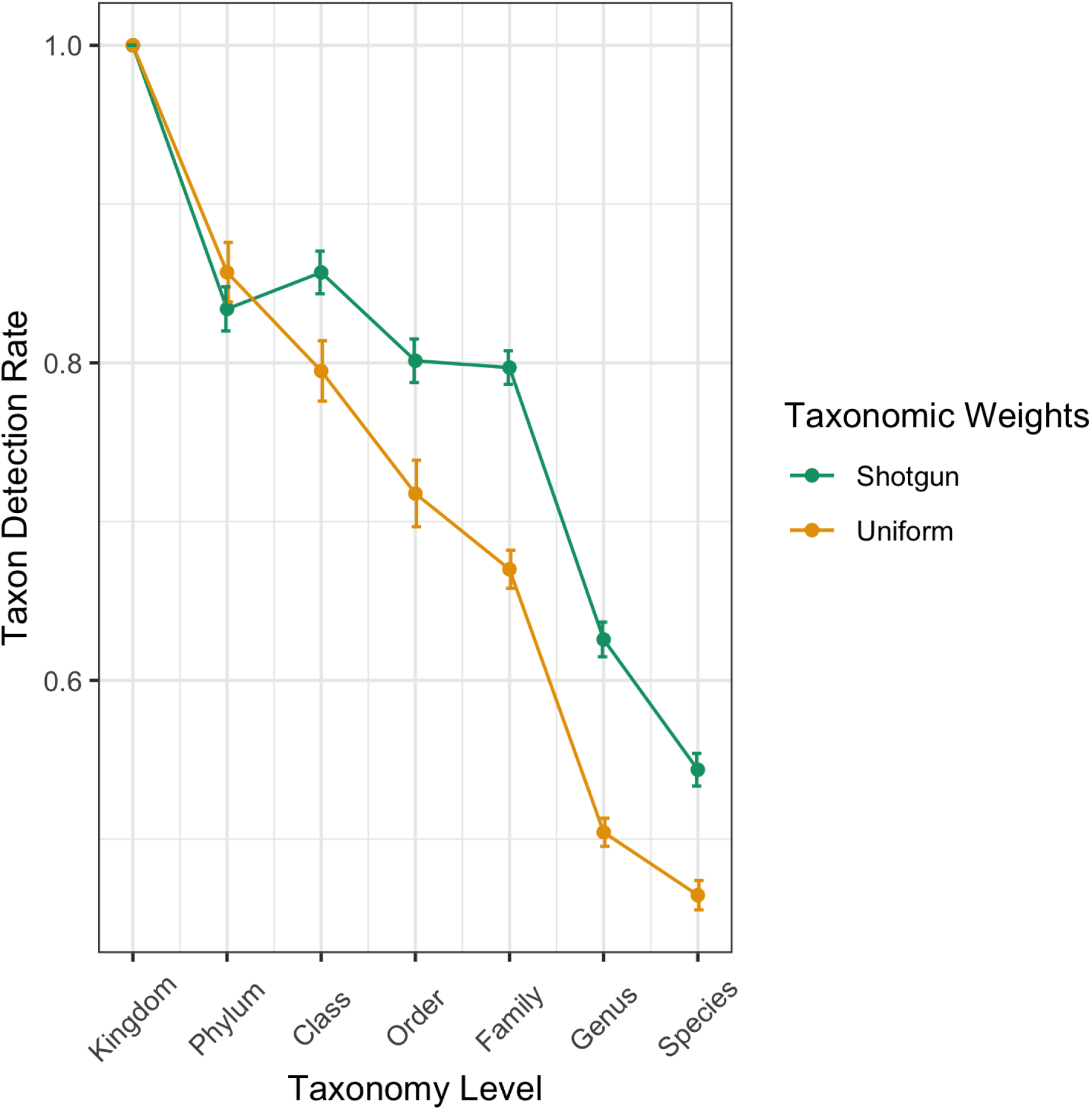
Taxonomic weights from shotgun sequencing improve agreement between amplicon and shotgun sequencing taxonomic compositions. Taxonomic weights derived from shotgun sequencing experiments make the taxa discovered in out-of-sample 16S sequencing samples agree more closely with shotgun sequencing experiments on the same samples. Taxon discovery rate (TDR) is the fraction of taxa detected using shotgun sequencing that were also found using 16S sequencing. Points show mean TDR across 71 stool samples for which paired 16S and shotgun sequencing exists. Error bars show standard errors across folds for 5-fold cross validation.

## References

1. Thompson, L. R. et al. en. Nature 551, 457–463 (2017).

2. Huttenhower, C. et al. Nature 486, 207 (2012).

3. Gonzalez, A. et al. Nat. Methods 15, 796–798 (2018).

4. Bokulich, N. A. et al. en. Microbiome 6, 90 (2018).

5. Cole, J., Konstantinidis, K, Farris, R. & Tiedje, J. Liu WT, Jansson JK (ed.) 515, 1–19 (2010).

6. Janda, J. M. & Abbott, S. L. en. J. Clin. Microbiol. 45, 2761–2764 (2007).

7. Jovel, J. et al. Front. Microbiol. 7 (2016).

8. Edgar, R. C. en. PeerJ 6, e4652 (2018).

9. Goodrich, J. K. et al. Cell 158, 250–262 (2014).

10. Bolyen, E. et al. PeerJ Prepr. 10.7287/peerj.preprints.27295v2 (2018).

11. Wang, Q., Garrity, G. M., Tiedje, J. M. & Cole, J. R. en. Appl. Environ. Microbiol 73, 5261–5267 (2007).

12. Rohwer, R. R., Hamilton, J. J., Newton, R. J. & McMahon, K. D. mSphere 3, e00327–18 (2018).

13. Tang, J., Iliev, I. D., Brown, J., Underhill, D. M. & Funari, V. A. J. Immunol. Methods 421, 112–121 (2015).

14. Ritari, J., Salojärvi, J., Lahti, L. & de Vos, W. M. BMC Genomics 16, 1056 (2015).

15. Fettweis, J. M. et al. BMC Genomics 13, S17 (2012).

16. Hain, T., Steinweg, C. & Chakraborty, T. J. Biotech. 126, 37–51 (2006).

17. Fuchs, T M., Eisenreich, W., Heesemann, J. & Goebel, W. FEMS Microbiol. Rev. 36, 435–462 (2012).

18. Thompson, F. L., lida, T. & Swings, J. Microbiol. Mol. Biol. Rev. 68, 403–431 (2004).

19. Fotedar, R et al. Clin. Microbiol. Rev. 20, 511–532 (2007).

20. Oliveira, A. et al. Front. Microbiol. 8, 1937 (2017).

21. Fouts, D. E. et al. PLoS Negl. Trop. Dis. 10, e0004403 (2016).

## References

22. Amir, A. et al. mSystems 2, e00191–16 (2017).

23. McDonald, D. et al. 2018.

24. Schulfer, A. F. et al. Nat Microbiol 3, 234–242 (2017).

25. Ruhe, J. et al. Front Plant Sci 7 (2016).

26. O’Brien, S. L. et al. Environ Microbiol 18, 2039–2051 (2016).

27. Lax, S. et al. Science 345, 1048–1052 (2014).

28. Zarraonaindia, I. et al. mBio 6 (2015).

29. Navas-Molina, J. A. et al. in Methods Enzymol 371–444 (2013).

30. Fang, X. et al. Front Microbiol 9 (2018).

31. Tripathi, A. et al. mSystems 3 (2018).

32. Delsuc, F. et al. Mol Ecol 23, 1301–1317 (2013).

33. Gibbons, S. M. et al. PLoS ONE 9, e97435 (2014).

34. Caporaso, J. G. et al. Genome Biol 12, R50 (2011).

35. Hyde, E. R. et al. mSystems 1 (2016).

36. Brazelton, W. J., Nelson, B. & Schrenk, M. O. Front Microbiol 2 (2012).

37. Vitaglione, P. et al. Am J Clin Nutr 101, 251–261 (2014).

38. Spirito, C. M., Marzilli, A. M. & Angenent, L. T. Environ Sci Technol 52, 13438–13447 (2018).

39. Pham, V. T. H. et al. Sci Rep 7 (2017).

40. McDonald, D. et al. ISME J. 6, 610 (2012).

41. Truong, D. T. et al. en. Nat. Methods 12, 902–903 (2015).

42. Callahan, B. J. et al. en. Nat. Methods 13, 581–583 (2016).

43. O’Leary, N. A. et al. en. Nucleic Acids Res. 44, D733–45 (2016).

44. Lemaître, G., Nogueira, F. & Aridas, C. K. J. Mach. Learn. Res. 18, 1–5 (2017).

45. Pedregosa, F et al. J. Mach. Learn. Res. 12, 2825–2830 (2011).

46. Bray, J. R. & Curtis, J. T. Ecol. Monogr. 27, 325–349 (1957).

## References

47. Segata, N. et al. Nat. Methods 9, 811 (2012).

